# Adolescent blockade of complement signaling in the lateral septum increases social novelty seeking behavior in male mice

**DOI:** 10.64898/2026.08.28.747845

**Authors:** Anthony Djerdjaj, William W. Taylor, Odette T. Amoakohene, Caroline J. Smith

## Abstract

Social behaviors are critical for survival and change dramatically over the lifespan. Adolescence is a critical period of development during which social novelty seeking peaks before declining into adulthood. Adolescence is also a time of pronounced neural circuit refinement as excess synapses are eliminated. One critical mechanism supporting this maturation of neural circuits is microglial pruning of synapses through the classical complement signaling cascade. However, it remains unclear how microglial pruning of the neural circuitry supporting social novelty preference shapes the trajectory of this behavior during adolescence. To address this, we blocked microglial complement-dependent pruning during adolescence by injecting neutrophil inhibitory factor (NIF; blocks the adhesion of ligands to CD11b/C3 receptor) in the lateral septum (LS), a key node in the social circuitry supporting social novelty preference, in male mice. We found that NIF administration into the LS during adolescence increased preference for the novel social chamber over the familiar as compared to control PBS administration. LS-NIF treatment had no impact on anxiety-like behavior in the light-dark box test and no effect on sociability. LS-NIF treatment also decreased the expression of immune-related genes in the LS as compared to PBS treatment. These data support the hypothesis that complement-dependent microglial synaptic elimination in the LS is critical for the developmental progression of social novelty preference.

## Introduction

Social behaviors are critical for survival across species and social impairments are hallmarks of many developmental neuropsychiatric disorders like autism spectrum disorder, schizophrenia, and anxiety disorders (Hamilton, 1964a, 1964b; Kennedy & Adolphs, 2012). Social behaviors undergo significant reshaping over the course of development to ensure age-appropriate expression of these behaviors. Adolescence is a critical period for social development during which peer social interactions and social novelty seeking peaks, supporting the transition from maternal dependence to adult social relationships (Spear et al., 2000).

Adolescence is also a period of dramatic brain remodeling via synaptic pruning and significant vulnerability for the onset of mental health disorders (Paus et al., 2008). In order to understand the developmental origins of adult social behavior and social dysfunction in disease, it is essential to fully elucidate the mechanisms by which social circuitry is established and matures during critical periods of social development.

The lateral septum (LS) is a key relay between cortical and subcortical structures and a critical component of social neural circuits (Menon et al., 2022). An array of LS-connections with regions like the bed nucleus of the stria terminalis, the amygdala, the ventral tegmental area, the paraventricular nucleus of the hypothalamus, and the ventral hippocampus have been shown to mediate social behaviors including aggression, affiliative behavior, and social memory in rodents (Mahadevia et al., 2021; Menon et al., 2022; Rigney et al., 2023, 2024; Wong et al., 2016). The LS also mediates social behaviors that are highly expressed during adolescence, like social play behavior and social novelty seeking. The LS densely expresses receptors for the neuropeptide vasopressin and blockade of these receptors has been shown to modulate social play during adolescence in a sex-specific manner, increasing it in males and decreasing it in females, in rats (Bredewold et al., 2014, 2025; Veenema et al., 2013). Social novelty preference, the tendency to prefer novel conspecifics over familiar ones, also peaks during adolescence and has been shown to be dependent on the LS. A recent study in mice demonstrated that inhibition of projections from the ventral hippocampus to the LS, and from these LS neurons to targets in the ventral tegmental area, disrupts social novelty preference and discrimination between familiar and novel conspecifics (Rashid et al., 2025). Research investigating the specific cells and signaling pathways in the LS involved in social behavior have revealed that deletion of oxytocin receptors or of tropomyosin receptor kinase B (TrkB), the receptor for brain-derived neurotrophic factor, in the LS of male mice significantly impairs social novelty preference (Mesic et al., 2015; Rodriguez et al., 2024).

Social novelty preference peaks during early adolescence, a period when social reward and risk taking are heightened, before declining as adolescence progresses into adulthood (Bian et al., 2022; Spear, 2000). Work has also demonstrated substantial molecular and synaptic changes in the LS during this period. Oxytocin receptor expressing cells in the LS are critical for social novelty preference and expression of this receptor similarly peaks in early adolescence and declines into adulthood (Bian et al., 2022; Hammock & Levitt, 2013; Mesic et al., 2015). This suggests that developmental changes in social novelty preference may be shaped by maturation of LS circuitry over the course of adolescence. However, little is known about the cellular mechanisms shaping LS circuits between adolescence and adulthood and how this contributes to LS dependent social behaviors like social novelty preference.

A key mechanism of adolescent neural circuit maturation is synaptic pruning. Synaptic pruning, the process of eliminating excess synapses, peaks first in early life and then again during adolescence (Faust et al., 2021; Riccomagno & Kolodkin, 2015). Pruning during adolescence results in a massive loss of synapses and a profound reorganization of the brain to support the transition to typical adult behaviors (Juraska & Drzewiecki, 2020; Spear, 2013). This process relies in part on microglia, the resident immune cells of the brain (Faust et al., 2021).

Microglia have been shown during critical periods of development to engulf and eliminate less active synapses (Schafer et al., 2012; Stevens et al., 2007). One mechanism by which these synapses are tagged and subsequently recognized by microglia for pruning is the classical complement cascade (Faust et al., 2021; Stevens et al., 2007). This involves the tagging of synapses with complement cascade components (C1q, C3) which are recognized by microglial complement receptor 3 (CR3), stimulating phagocytosis of the synapse (Faust et al., 2021; Nandi et al., 2025). Complement-dependent phagocytosis by microglia has been shown to be critical for the development of adolescent social behaviors like social play, such that a function-blocking antibody against CD11b (a component of CR3) in early life reduces the development of play behavior in adolescence (VanRyzin et al., 2019). Furthermore, blocking complement-dependent microglial pruning in the nucleus accumbens during adolescence using neutrophil inhibitory factor (NIF), a high-affinity CD11b antagonist that stops CR3 formation, attenuated the developmental decline of social play and increased social interaction with a familiar conspecific (Kirkland et al., 2024; Kopec et al., 2018; Muchowski et al., 1994). Social novelty preference during adolescence is also shaped by microglia pruning. Genetic mouse models of autism have demonstrated impaired social novelty preference that is associated with disrupted microglial function and can be rescued by modulating microglial pruning (Ju et al., 2025; Lian et al., 2026). Despite this evidence showing the importance of microglial pruning to social development and our understanding that the LS is critical for adolescent social behaviors, no studies have investigated the importance of adolescent microglial pruning specifically in the LS to the expression social novelty preference.

In this study, we examined the impact of blocking microglial complement-dependent pruning in the LS during adolescence on social novelty preference, sociability, and anxiety-like behavior. To accomplish this, we bilaterally injected NIF or PBS control into the LS of adolescent male mice at postnatal day (PN)30-31, a time point at which local NIF administration has previously been shown to block pruning in the nucleus accumbens (Kopec et al., 2018). We assayed the impact of this manipulation on behavior between PN38-42, when adolescent social behaviors have typically declined (Kopec et al. 2018) using the light-dark box test, three-chamber sociability and social novelty preference test, and free roaming social interaction test. We then isolated RNA from LS punches to determine the transcriptional consequences of NIF injection. Given evidence that declining social novelty preference coincides with synaptic elimination over the course of adolescence and the importance of microglial pruning to the trajectory of other adolescent behaviors like social play, we hypothesized that blocking complement-dependent microglial pruning in the LS during adolescence would increase social novelty preference.

## Materials and Methods

### Animals

Male C57BL/6 mice were purchased from Jackson Laboratory (#000664) and allowed to acclimate to the vivarium in the Boston College Animal Facility for 5 days before any procedures were carried out. Mice arrived at postnatal day 24 (P24) and were housed 4 to a cage. The vivarium was maintained on a 12h light/dark cycle (6am-6pm) and food and water were available *ad libitum*. All procedures were approved by the Boston College Institution Animal Care and Use Committee and adhered to the Public Health Service *Guide for the Care and Use of Laboratory Animals*.

### Surgical Procedures

To determine the effects of blocking microglial complement-dependent synaptic pruning within the LS during adolescence, neutrophil inhibitory factor (NIF, R&D Systems) or sterile phosphate buffered saline (PBS) were microinjected into the ventral portion of the LS (LSv) of 13 male mice (7 NIF-treated mice, 6 PBS-treated mice) at PN30-31. We chose to target the ventral LS as this is the portion of the LS where oxytocin receptor density (important for social novelty seeking) is higher during adolescence than in adulthood in rodents (Smith et al., 2017). NIF was prepared in sterile PBS according to the manufacturer’s recommendations at a concentration of 200ng/uL. This drug choice, timing, and dose were chosen based on previous work showing that local NIF transfusion in the nucleus accumbens at PN30 prevents C3-C3R interactions and increases later social play behavior in male rodents. On PN30-31, experimental mice underwent surgery under inhaled anesthesia (0.5-3% v/v isoflurane in O_2_). At the start of the surgical procedure, mice received a subcutaneous Ketofen injection (5mg/kg; Covetrus). Sterile PBS or NIF was deposited bilaterally into the LSv (from bregma: A/P: +0.75; M/L: +/-0.55, D/V: -3.65) at a rate of 100nL/min to a total volume of 300nL. 5 minutes were allowed for diffusion. Wound clips were used to seal the site of incision and mice were allowed to recover on a heating pad. Post-operative care was conducted for 4 days to ensure animals’ well-being and recovery. This included daily body weight measurement, assessment of body posture and piloerection, and inspection of the incision site for wound healing. Animals were allowed to rest and recover for ∼ 1 week prior to behavioral testing (see **Fig. 1A** for experimental timeline).

**Figure 1.**
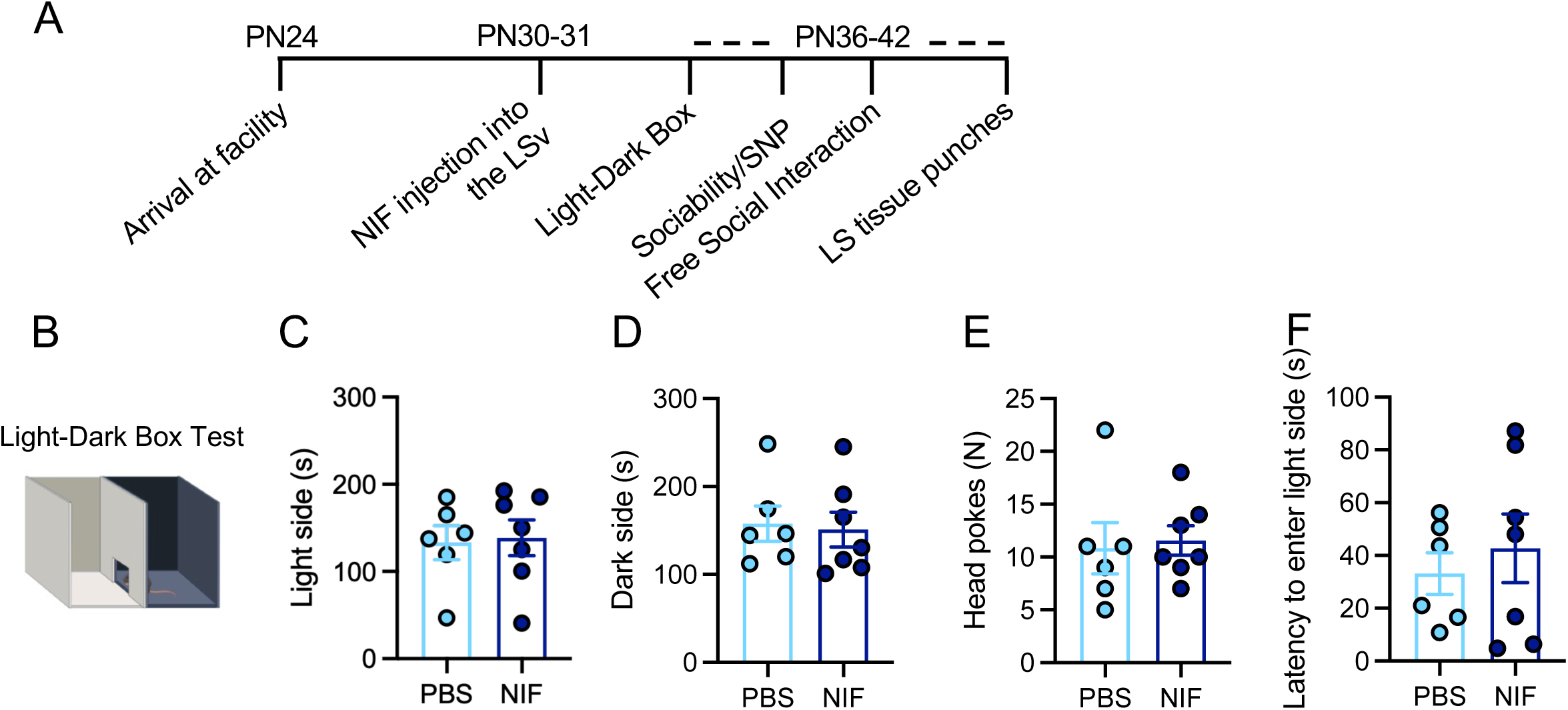
**A**) Experimental timeline. Animals arrived at P24 to our animal facility. At P30/31 animals underwent stereotaxic surgery to microinject NIF into the LSv. At P36-42 animals underwent a series of behavior assays including Light-Dark box, Sociability, Social Novelty Preference and Free Social Interaction. After this, tissue punches were collected from the LS to assess gene expression. **B**) Light-Dark Box apparatus **C-F**) There were no significant effects of NIF treatment on time spent in the light (**C**), time spent in the dark (**D**), head pokes (**E**), or latency to enter the light side (**F**). Data represent mean +/-SEM, un-paired T-test results. PN= postnatal day, PBS= phosphate-buffered saline, NIF= neutrophil inhibitory factor.

### Behavioral Testing

Behavior testing took place from PN36-42 during the second half of the light phase (afternoon). Animals were handled daily in the 3 days prior to behavior and habituated to the testing room and testing apparatus prior to testing.

#### Light/Dark Box

To assess ‘anxiety-like’ behavior, mice underwent the light/dark box test. Mice were placed in the testing room one hour prior to testing. At the start of each test, experimental mice were placed into the dark side of a two-chamber arena and allowed to freely explore each chamber over the course of 5 minutes. Each test was recorded and behavioral videos were scored using Solomon Coder by an experimenter blind to the treatment condition of each subject. The amount of time mice spent on each side of the arena (seconds), the number of head pokes into the light side (N), and the latency to enter the light side (seconds) were quantified.

#### Sociability and Social Novelty Preference

To assess sociability (preference to investigate a novel social stimulus vs. a novel object) and social novelty preference (preference to investigate a novel social vs. a familiar social stimulus), a three-chambered test was used.

##### Habituation

The day prior to testing, mice were habituated to the testing environment. Experimental mice were placed into the middle chamber of a three-chamber arena with two empty stimulus containers (Plexiglass rod sides) in the left and right chambers. Mice were allowed to explore the arena for 5 minutes. Separately, target conspecifics were placed into the stimulus containers for 5 minutes each to allow for acclimation to the container.

##### Sociability

For sociability tests, a novel object (rubber duck) and a novel sex and age matched social stimulus were each confined to a container and placed in either the left or right chamber (counterbalanced across tests). Experimental mice were then placed into the middle empty chamber and allowed to freely move between chambers and investigate each stimulus over the course of 5 minutes. All tests were recorded and videos were scored by a blinded observer using Solomon Coder. Time spent in each of the 3 chambers (seconds), time spent investigating the object or social stimulus (direct sniffing between the bars of the containers; seconds) and time spent climbing each container (seconds) were quantified. Preference for the social chamber was calculated by dividing the time spent in the social chamber by the total time spent in either the social or the object chamber and multiplying by 100 to quantify a percentage.

##### Social Novelty Preference

After the 5-minute sociability test, a social novelty preference test was conducted. The container holding the novel object was removed and replaced with a novel sex- and age-matched social stimulus confined to a container. Experimental mice were allowed to investigate each stimulus over the course of 5 minutes. All tests were recorded and videos were scored by a blinded observer using Solomon Coder. Time spent in each of the 3 chambers (seconds), time spent investigating each social stimulus (direct sniffing between the bars of the containers; seconds) and time spent climbing each container were quantified. Preference for the novel chamber was calculated by dividing the time spent in the novel chamber by the total time spent in either the novel or the familiar chamber and multiplying by 100 to quantify a percentage.

#### Free-roaming Social Exploration

To assess more exploratory social behaviors (sniffing, pinning, allogrooming), a one-on-one social exploration test was performed. Each experimental mouse was placed into a standard housing cage with beta chip bedding and a wire lid 1 h prior to testing. Testing consisted of a novel same-sex conspecific being introduced into the experimental mouse’s cage for 5 minutes. All tests were recorded and scored using Solomon Coder. Exploratory social behaviors that were quantified included general investigation (seconds), anogenital sniffing (seconds), and allogrooming (seconds). Non-social behaviors that were quantified included general cage exploration (seconds), rearing (seconds), digging (seconds), and autogrooming (seconds).

### Tissue Collection and punches

All animals were euthanized using CO_2_ inhalation following the completion of behavioral testing. Brains were extracted, flash frozen in 2-methylbutane over dry ice, and stored at -80°C until sectioning. Punches were collected using a 1 mm diameter core sampling tool. Brains were mounted in a sterilized cryostat (thoroughly cleaned with 70% ethanol between each sample).

Brains were sliced until about A/P +0.97 from bregma and punches of the ventral portion of the lateral septum were collected by inserting the sterilized core sampling tool to a depth of 1 mm. Punches were placed into microcentrifuge tubes and immediately frozen at -80°C until RNA extraction.

### RNA Extraction

Frozen samples were homogenized in 500mL of TRIzol Reagent (Sigma-Aldrich) using a Tissue Tearer and vortexed on a MixMate at 2000rpm for 10 min. 100mL of chloroform (Sigma-Aldrich) was added to each tube and vortexed at 2000rpm for an additional 2 min. Samples were allowed to phase separate before being centrifuged at 11,800rpm for 15 min at 4°C. The clear aqueous layer was collected into a fresh tube and 200mL of isopropanol (Sigma-Aldrich) and 2mL of glycogen (Invitrogen) were added to precipitate and visualize RNA. Samples were vortexed at 2000rpm for 1 min and incubated at room temperature for 10 min before being centrifuged at 11,800rpm for 10min at 4°C. Each tube was checked for a pellet and supernatant was discarded quickly, ensuring that the pellet was not lost. Samples were then rinsed twice with ethanol. 500mL of cold 75% EtOH was added to each sample and samples were centrifuged at 9000rpm for 10 min at 4°C. EtOH was decanted and this process was repeated. Following the second EtOH rinse, a q-tip was used to remove excess EtOH from each tube and samples were allowed to dry for ∼12 minutes. Finally, the pellets were suspended in 10mL of nuclease-free water (Thermofisher Scientific) and immediately frozen at -80°C until cDNA synthesis.

### cDNA synthesis

Before cDNA synthesis, RNA samples were nanodropped to determine concentration and purity of each sample. Based on observed concentrations, samples were diluted to a 1000ng/uL concentration in nuclease free water. cDNA was then synthesized using the QuantiTect Reverse Transcription Kit (Quiagen). Briefly, RNA was pre-treated with 2mL of gDNase for 2 minutes at 42°C in the thermocycler to remove genomic DNA contamination. A master mix containing buffer, primer mix, and reverse transcriptase was added to each sample. Samples were heated to 42°C for 30 min and 95°C for 3 minutes in the thermocycler. Once completed, 80mL of NF water was added to each sample and samples were divided between 5 tubes for a final concentration of 200ng/mL of RNA.

### qPCR

Quantitative real-time PCR (qPCR) was conducted using QuantiTect SYBR Green PCR kit (Quiagen) on a QuantStudio 3 real-time PCR machine (Thermofisher Scientific). All samples were run in duplicate. Gene expression was analyzed for the following genes: *Cd11b*, *Cd68*, *Tlr4*, and *IL-1*β. 18S was used as a house-keeping gene and relative gene expression was calculated using the 2^-ΔΔCt^ method, relative to *18S* and the lowest sample on the plate (Livak & Schmittgen, 2001; Williamson et al., 2011). If samples showed multiple melt curve peaks (indicative of contamination) or if duplicate values differed by >1 fold change, they were removed from analyses prior to unblinding. Primer sequences are as follows: *18S*: Forward (F): GAATAATGGAATAGGACCGC, Reverse (R:) CTTTCGCTCTGGTCCGTCTT *Cd11b*: F:CTATTTGTTCGGCTCCAAC R:GCATCAAAGAGAACAAGG, *Cd68*: F:CCCACCTGTCTCTCTCATTTC R:GTATTCCACCGCCATGTAGT *Tlr4:* F:CAGCAGAGGAGAAAGCAT R:CACCAGGAATAAAGTCTCTG *IL-1*β: F:GCATCAAAGAGAACAAGG R:CACAGGCTCTCTTTGAAC.

### Statistics

Statistical analyses were conducted using Graphpad Prism version 11 software. Un-paired t-tests were used to compare the effects of PBS vs. NIF treatment on all behavioral measures and gene expression analyses. Significance was set at p<0.05.

## Results

### LS-NIF injection has no impact on Light-Dark Box behavior

There were no significant effects of NIF treatment compared to PBS control on time spent in the light side of the light-dark box (**Fig. 1B-C**; t_(11)_=0.197, p=0.848), time spent on the dark side (**Fig. 1D**; t_(11)_=0.233, p=0.820), head pokes into the light side of the box (**Fig. 1E**; t_(11)_=0.274, p=0.789), or latency to enter the light side of the box (**Fig. 1F**; t_(11)_=0.607, p=0.556).

### LS-NIF injection increases preference for the novel chamber in the social novelty preference test, but does not alter sociability or free-roaming social interaction

In the sociability assay (**Fig. 2A**), there were no significant effects of NIF treatment compared to PBS control on any behavioral outcomes including % social chamber time (**Fig. 2B**; t_(11)_=0.531, p=0.606), social chamber time (**Fig. 2C**; t_(11)_=0.440, p=0.669), object chamber time (**Fig. 2D**; t_(11)_=0.666, p=0.519), middle chamber time (**Fig. 2E**; t_(11)_=1.21, p=0.251), social investigation (**Fig. 2F**; t_(11)_=0.511, p=0.620), object investigation (**Fig. 2G**; t_(11)_=0.594, p=0.564), social climbing (**Fig. 2H**; t_(11)_=1.09, p=0.310), and object climbing (**Fig. 2I**; t_(11)_=1.15, p=0.273).

**Figure 2.**
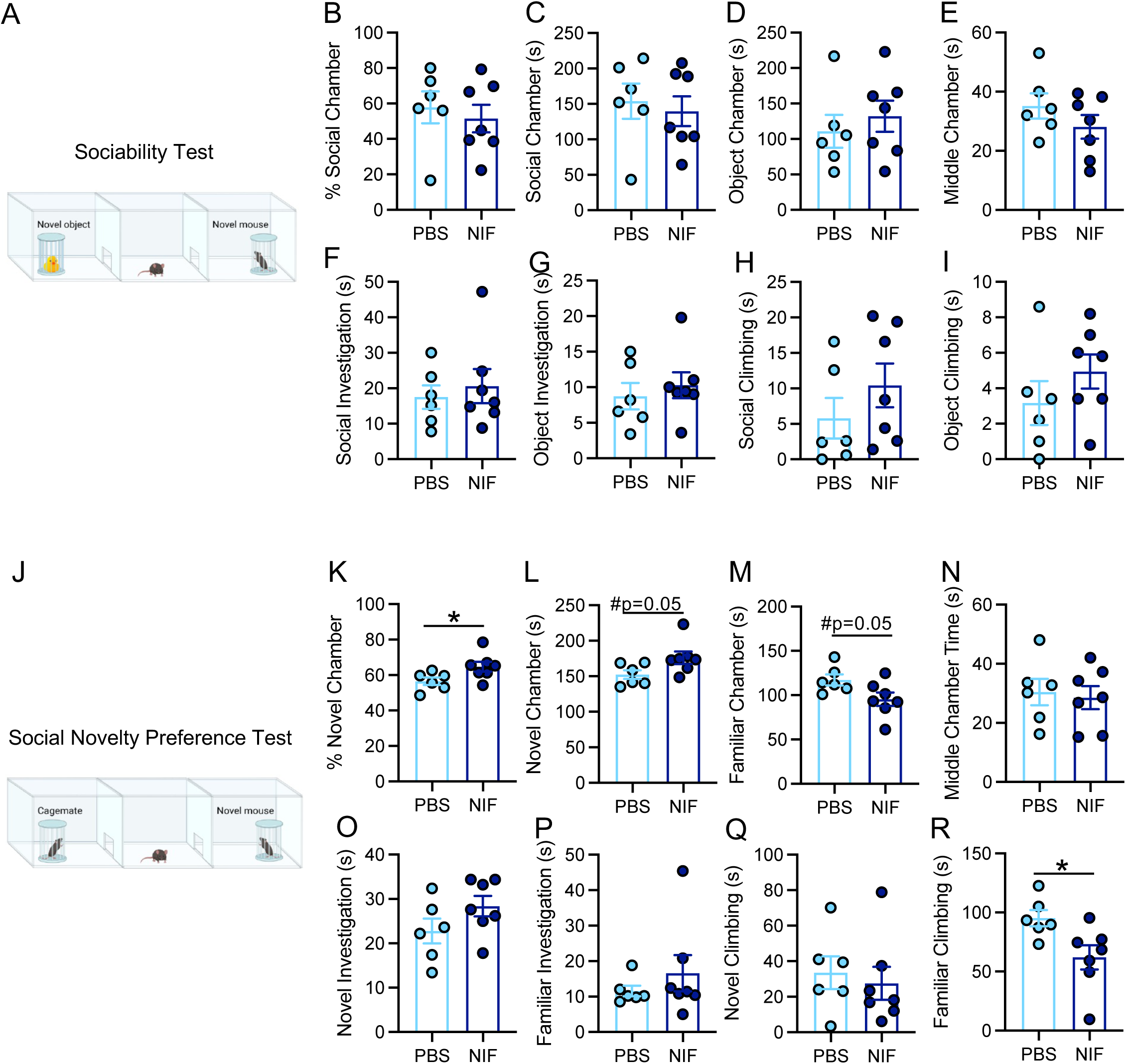
**A**) Schematic representing the sociability assay. **B-E**) Chamber results of the sociability assay. NIF treatment had no effect on % social chamber time (**B**), social chamber time (**C**), object chamber time (**D**), or middle chamber time (**E**). Similarly, NIF treatment had no effect on social investigation (**F**), object investigation (**G**), social climbing (**H**), or object climbing **(I**). **J**) Schematic representing the social novelty preference test. **K-N**) Chamber results of the social novelty preference test. NIF treatment significantly increased % preference for the novel chamber as compared to control (**K**). It tended to increase novel chamber time (**L**) and decrease familiar chamber time (M) but had no impact on middle chamber time (N). There was no effect of NIF treatment on novel investigation (**O**), familiar investigation (**P**), or novel climbing (**Q**), but a significant decrease in familiar climbing (**R**). Data represent mean +/-SEM, un-paired t-test results. *p<0.05. #p<0.07. PBS= phosphate-buffered saline, NIF= neutrophil inhibitory factor.

However, in the social novelty preference test (**Fig. 2J**), there was a significant increase in the preference for the novel chamber following NIF treatment as compared to PBS control (**Fig. 2K**; t_(11)_=2.30, p=0.042). This was driven by trends towards increased novel chamber time (**Fig. 2L**; t_(11)_=0.2.13, p=0.057) and decreased familiar chamber time (**Fig. 2M**; t_(11)_=2.18, p=0.052). There was no effect of NIF treatment on middle chamber time in the social novelty preference test (**Fig. 2N**; t_(11)_=0.325, p=0.751), nor was there an effect on novel investigation (**Fig. 2O**; t_(11)_=0.1.56, p=0.147), familiar investigation (**Fig. 2P**; t_(11)_=0.873, p=0.402), or novel climbing (**Fig. 2Q**; t_(11)_=0.454, p=0.659). There was a significant decrease in time spent climbing the familiar chamber following NIF treatment as compared to PBS control (**Fig. 2R**; t_(11)_=2.58, p=0.026).

In the free-roaming social interaction test, no significant differences were found between NIF and PBS treated animals (see **Table 1** for complete means and statistics).

**Table 1.**
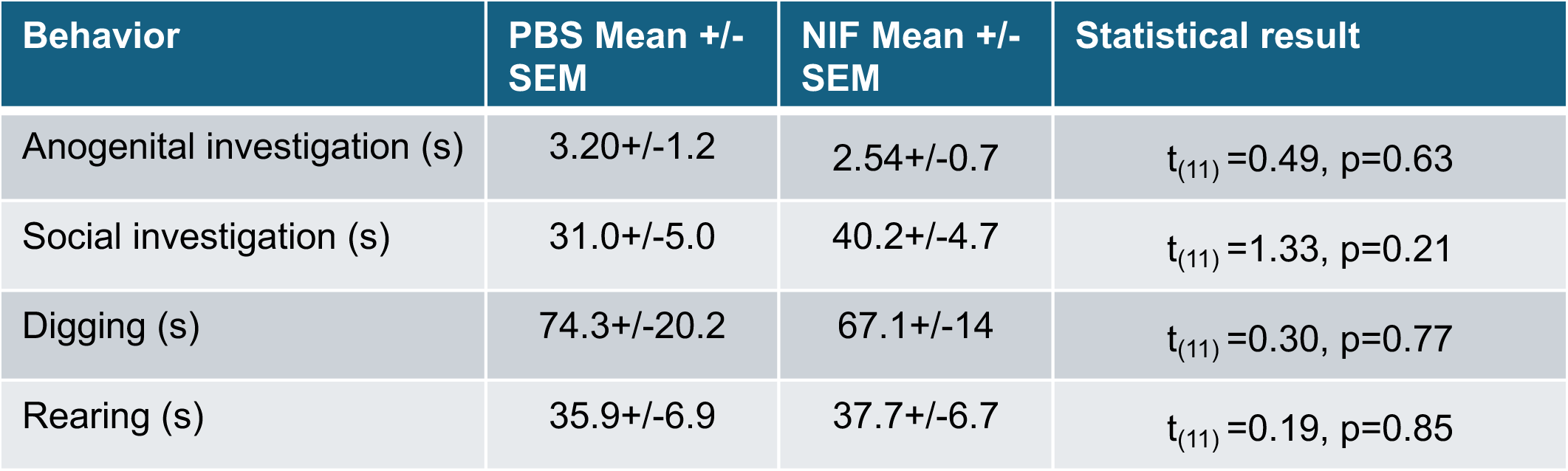
Behavior in free-roaming social interaction test. Un-paired t-test results of PBS vs NIF treatment on the below outcome measures. (s): seconds. PBS= phosphate buffered saline, NIF= neutrophil inhibitory factor.

### LS-NIF injection decreases immune gene expression in the LS

We collected tissue punches from the LS (**Fig. 3A**) after either NIF or PBS treatment to determine how NIF administration impacted immune markers in the LS. NIF treatment significantly decreased the mRNA expression of *Cd11b* (a protein subunit of C3R which is the target of NIF blockade; **Fig. 3B,** t_(9)_=2.37, p=0.042) as compared to PBS control. There was also a significant decrease in *IL-1*β (**Fig. 3B,** t_(7))_=2.40, p=0.048), and a trend towards a decrease in *Tlr4* mRNA expression (**Fig. 3B,** t_(7)_=1.92, p=0.096). There was no significant difference in *Cd68* (**Fig. 3B,** t_(9))_=1.64, p=0.136).

**Figure 3.**
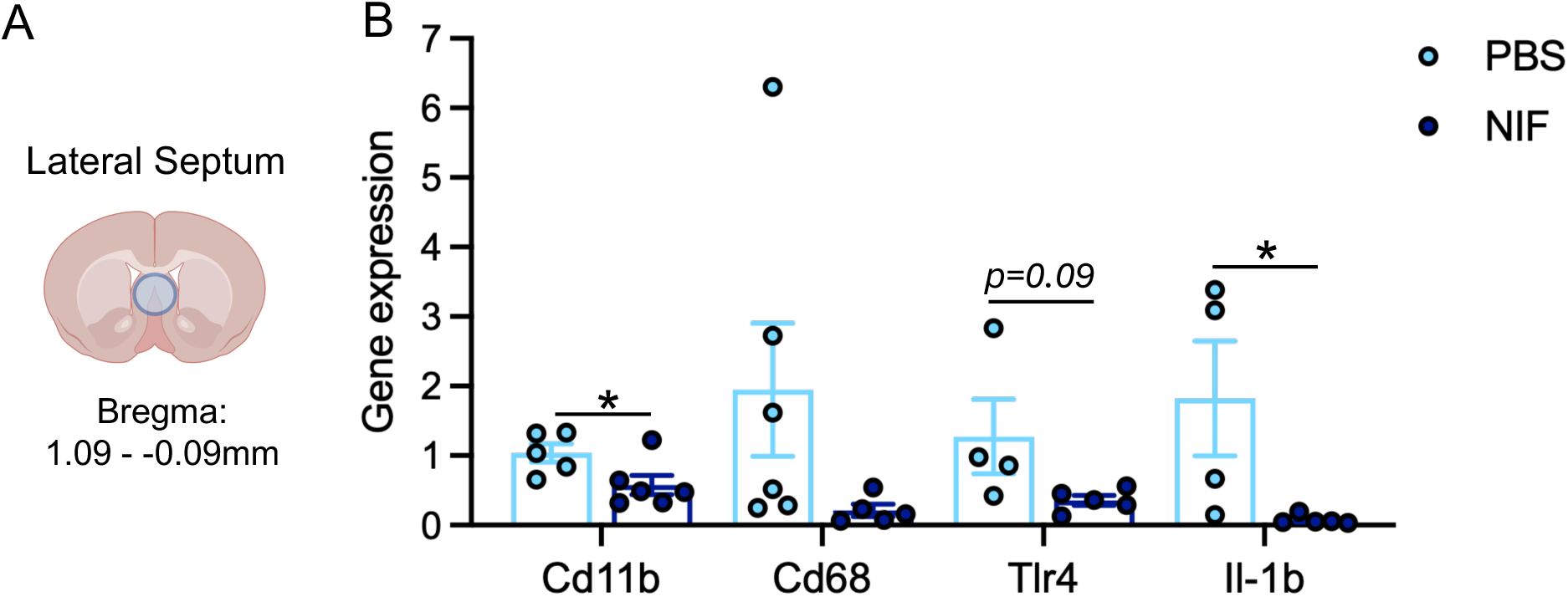
**A**) Schematic representing location of tissue punches in the LS. **B**) Gene expression data relative to *18S* and the average of the PBS control group for each gene of interest. NIF treatment decreased mRNA expression for *Cd11b* and *Il-1*β in the LS as compared to control and tended to decrease *Tlr4* mRNA expression. Data represent mean +/-SEM, un-paired t-test results. *p<0.05. #p<0.07. PBS= phosphate-buffered saline, NIF= neutrophil inhibitory factor.

## Discussion

Overall, we found that adolescent NIF-mediated blockade of microglial complement-signaling in the LS during adolescence increased subsequent preference for the novel social chamber over the familiar chamber as compared to control. To the best of our knowledge, this work is the first to investigate the role of microglia in the shaping of social behavior in the LS. This finding is consistent with the idea that blockade of complement-dependent synaptic pruning during a critical period for this process prevents typical declines in social novelty seeking. LS-NIF treatment had no impact on anxiety-like behavior as measured by the light-dark box test and no effect on overall sociability. Furthermore, LS-NIF treatment decreased the expression of immune-related genes in the LS, suggesting that in addition to decreasing the function of C3R/CD11b, NIF may also decrease other inflammatory pathways in microglia as well. Together, these data suggest, for the first time, that complement-dependent microglial synaptic elimination in the LS is critical for the developmental progression of social novelty preference.

Social novelty seeking behavior declines between adolescence and adulthood and is mediated, in part, by the LS. At the same time, the brain undergoes significant synaptic pruning. Therefore, we hypothesized that developmental synaptic pruning in the LS might be responsible for declines in social novelty seeking between these two periods. Consistent with this hypothesis, we find that blockade of complement signaling locally in the LS during early adolescence is sufficient to increase preference for a chamber containing a novel social conspecific during late adolescence when declines in other adolescent specific social behaviors such as social play have previously been observed. These findings are consistent with previous work in other social brain regions, such as the nucleus accumbens (Kopec et al., 2018) and medial amygdala (VanRyzin et al., 2019) showing that microglial synapse elimination is essential to both the establishment of and then decline in adolescent social play behavior (Nelson & Lenz, 2017). Interestingly, a previous study of NIF administration also observed specific effects on social novelty preference but not sociability more broadly. Kirkland et al. (2024), found that NIF administration into the nucleus accumbens at P30 of male rats led to increased *familiar* investigation (as a percentage of familiar chamber time) in adulthood. This is in contrast to the increase in *novelty* preference observed in our experiment. This suggests that the LS and the nucleus accumbens may have distinct roles in the investigation of, and preference for, novel and familiar social stimuli.

The LS also plays an important role in the circuits that support fear and anxiety-like behaviors in rodents. Many types of stress activate the LS and modulating the activity of stress responsive cells in the LS can modify the expression of fear and anxiety-like behaviors (Anthony et al., 2014; Rizzi-Wise & Wang, 2021; Sheehan et al., 2004). For example, a subset of stress-sensitive LS neurons express receptors for corticocotropin-releasing hormone (CRH) and project to the hypothalamus to promote anxiety-like behaviors in response to stress (Anthony et al., 2014). Optogenetic activation of these cells drives a persistent anxiogenic effect in the light-dark box, open field, and novel object tests, while optogenetic inhibition of these cells does the opposite (Anthony et al., 2014). Despite this, we found no effects of LS-NIF treatment on anxiety-like behavior in the light-dark box suggesting that LS microglial complement signaling does not play a role in shaping adolescent anxiety-like behavior within the context of the LS. Interestingly, microglia have been shown to modulate anxiety-like behaviors by engulfment of GABAergic dendritic spines in the central amygdala (Chen et al., 2024). This suggests that the involvement of microglia in the organization of anxiety circuits is likely centered in other brain regions or not organized during the developmental window assessed in this study.

Our results show that preventing microglial complement signaling in the LS during adolescence heightens social novelty seeking behavior. One outstanding question is what the cellular or sub-circuit mediators of this effect might be within the LS. The LS is a neurochemically diverse structure containing glutamatergic, GABAergic, and cholinergic neurons as well as receptors for an array of peptides and neurotransmitters including vasopressin, oxytocin, corticotropin releasing hormone, estrogen, and dopamine (Isaac et al., 2025; Menon et al., 2022; Swanson & Risold, 2000). Among these molecular players in the LS, oxytocin has been implicated as key to social novelty preference. LS oxytocin receptor (OTR) knockout was shown to significantly impair preference for social novelty, while activation of OTR+ cells in the LS ameliorated SNP deficits observed in mouse models of autism (Horiai et al., 2020; Mesic et al., 2015). Expression of OTRs in the LS also peaks in early adolescence and decreases into adulthood, mirroring the progression of social novelty preference (Hammock & Levitt, 2013). While the exact mechanism mediating OTR decline in the LS during adolescence is unknown, it is possible that microglial pruning of OTR+ synapses plays a role in the developmental progress of social preferences. This is an important area for further research.

Our data also demonstrate transcriptional changes in the LS following NIF injections. NIF binds CD11b, blocking complement signaling, which has been shown in rodent stroke models to reduce ischemic injury and inflammation (Jiang et al., 1998; Mackay et al., 1996; L. Zhang et al., 2003). While a reduction in inflammatory markers signals that transcriptional changes are taking place, no studies to our knowledge have directly investigated the transcriptional consequences of NIF administration in the central nervous system. Our findings that CD11b and IL1b are significantly downregulated following NIF suggest that not only is complement signaling disrupted, but that the inflammatory profile of microglia is changed. IL1b is a marker for microglia and its upregulation is a key marker of inflammation and shifting microglia morphology in the brian (Liu & Quan, 2018). While limited, our results suggest that microglia could be changing their inflammatory profile more broadly in response to NIF than theorized.

This study has several important limitations. While this is the first study to examine the role of microglia in shaping LS circuits during development, it is a small study, and future investigations should aim to expand upon these findings. Second, we limited our investigation to males. We made this decision because the role of microglial pruning during adolescence is better defined in males and the time course is better established (Kopec et al., 2018). However, there are known sex differences in microglial sculpting of social circuits and the role of microglia in organizing female social circuits is critical to better understand (Bordt et al., 2020; VanRyzin et al., 2019, 2020). Future work should aim to determine whether microglial complement signaling plays a role in the LS in females as well. This is especially important given that the LS is a brain region replete with sex differences in cellular and molecular functions relevant to social behavior (Bredewold et al., 2025; Rigney et al., 2024). Finally, our NIF manipulation is not pharmacologically specific to microglia. However, while NIF blocks C3-C3R interactions regardless of cell type, the C3 receptor is almost exclusively expressed by microglia in the mouse and human brain, with only very low expression in oligodendrocyte precursor cells, and CD11b is a constitutive marker for microglia (Y. Zhang et al., 2014). Therefore, it is highly likely that the effects observed here are dependent on microglial function specifically.

In summary, this work suggests that complement-dependent microglial synaptic pruning in the LS is a key mechanism supporting the developmental progression of social novelty preference. While more work is required to determine the downstream molecular targets impacted by disrupted microglial pruning, these studies are an important step towards elucidating the neurobiology supporting the development of social behaviors in adolescence.

## Ethics and Integrity

All data will be publicly available through the Boston College Dataverse. Research in the Smith lab is supported by the NIEHS (Grant No. R00ES033278) and the Department of Psychology and Neuroscience at Boston College. The authors have no conflicts of interest to disclose and all experiments were approved by the Boston College Institutional Animal Care and Use Committee.

## Acknowledgments

We are grateful for animal husbandry and care provided by the Animal Care Facility at Boston College.

## Notes

### Competing Interest Statement

The authors have declared no competing interest.

